# *Semecarpus anacardium* Linn. leaf extract exhibits activities against breast cancer and prolongs the survival of tumor-bearing mice

**DOI:** 10.1101/2020.01.08.898940

**Authors:** Rajesh Kumar Singh, Bhagaban Mallik, Amit Ranjan, Ruchita Tripathi, Sumit Singh Verma, Vinamra Sharma, Subash Chandra Gupta, Anil Kumar Singh

## Abstract

*Semecarpus anacardium* Linn. is commonly used in various traditional medicines from ancient times. The nuts have been described in Ayurveda medication systems to treat numerous clinical ailments. However, isolating phytochemical constituents from nuts remains challenging and exhibits cytotoxic effects on other cells. In this study, we have standardized procedures for isolating phytochemicals from the leaf extract. The ethyl acetate leaf extract selectively affects cancer cells in a dose-dependent manner (IC50: 0.57 μg/ml in MCF-7 cells) in various cancer cell lines.

Next, we examined if the extract incubation could induce cell cycle arrest and suppress cell migration in the cell culture model. Consistent with this idea, the leaf extract could potentially affect the aggressive migration nature of cancer cells. Moreover, oral administration of extract significantly restored tumor growth in mice. Together, these observations suggest the anti-cancer activities of *S. anacardium* leaf potential for both in vitro and in vivo models.

**Graphical Abstract:** 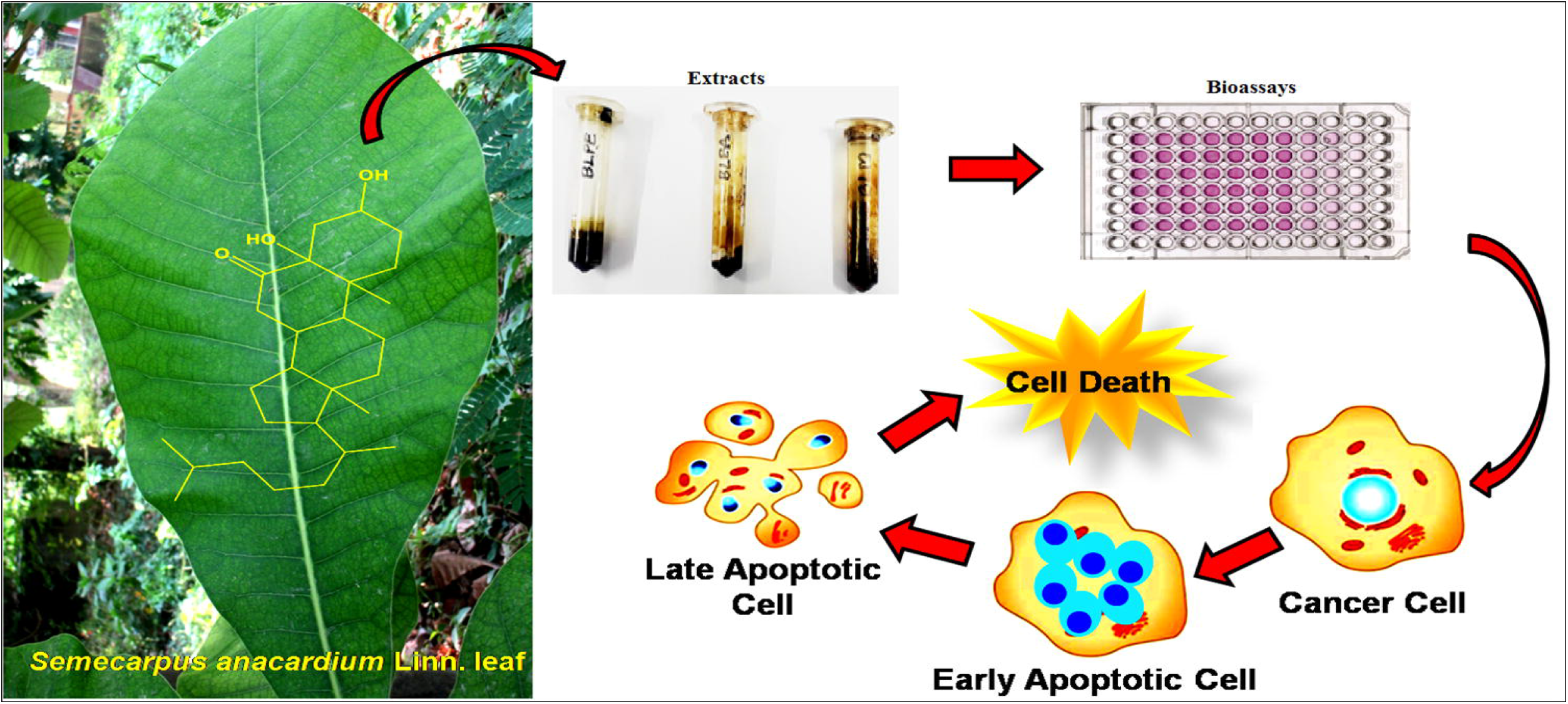

## 1. Introduction

Breast Cancer is one of the leading diseases among women and creates a significant global disruption in the healthcare system. In 2018, 2.09 million new breast cancer cases were diagnosed, with 0.63 million mortality; however, a recent survey suggests a projection of 3.06 million new cases and 0.99 million deaths by 2040 (Bray *et al*. 2018). Though much progress has been made in designing therapeutics, however; the treatment of breast cancer remains at its infant stage. Moreover, chemotherapy remains the common pharmacological approach for its treatment with many limitations, including severe toxic side effects leading to death in cancer patients (Arya *et al*. 2015). Additionally, many drugs designed for breast cancer are often cytotoxic for normal cells and induce cell death by cell cycle arrest, inhibition of DNA synthesis, necrosis, or apoptosis. The adverse effects of these drugs cause severe damage to the vital organs and normal proliferating cells such as the heart, liver, kidneys, hair follicles, stem cells, etc. (Vijayalakshmi *et al*. 1997; Vijayalakshmi *et al*. 2000). Moreover, new efficient and non-toxic chemotherapeutic agents need to be discovered to overcome this consequence.

Nevertheless, natural products have been used to treat many clinical ailments as described in different traditional systems of medicine worldwide since antiquity (Kim *et al*. 2018). Among these, Ayurveda is one of the traditional medicine systems prevalent in the Indian subcontinent, which describes the holistic approach to treating various human clinical ailments, including processed medicinal plants, some animal parts, and minerals (Joshi *et al*. 2010). Therefore, medicinal plants described in Ayurveda might serve as a potential source for rational drug design for many diseases, including cancer, neurological disorders, and lung and liver diseases. Many phytoconstituents from medicinal plants contain potential bioactive molecules such as alkaloids from *Rauwolfia serpentina, Holarrhena antidysenterica, Piper longum, Withania somnifera, Curcuma longa*, etc. for the treatment of hypertension, Amoebiasis, cough, stress, and cancer, etc. (Patwardhan and Vaidya 2010). Several compounds isolated from Azadirachta indica and *Premna herbacea* modulate numerous signaling pathways in the cell and show potent anti-cancer activity (Awasthee *et al*. 2018; Gupta *et al*. 2017). For instance, many bioactive compounds, including curcumin, flavopiridol, genistein, capsaicin, berberine, boswellic acid, caffeic acid, resveratrol, etc., are in a clinical trials for the treatment of cancer (Pandey *et al*. 2017).

Additionally, *Semecarpus anacardium* Linn. (Family: Anacardiaceae) is described in Ayurveda for treating several clinical ailments such as inflammation, vitiligo, neurological disorders, microbial infection, geriatric problem, diabetes, cancer and baldness, etc. (Mathivadhani *et al*. 2007b). Besides, *Semecarpus anacardium* Linn. fruits have been used in numerous Ayurvedic formulations. However, recent studies have indicated it as a potential source of antioxidant, antimicrobial, hypoglycemic, anti-inflammatory, anti-cancerous, antiatherogenic and stimulant for the central nervous system, skin diseases, and promoters for hair growth (Semalty *et al*. 2010). In contrast, the nut exhibits cytotoxicity activity against breast, liver, lung, cervical, and blood cancer cells (Selvam *et al*. 2004; Semalty *et al*. 2010; Sharma *et al*. 1995). Besides, the anti-tumor activity has also been evaluated in tumor-bearing animal models. However, isolating phytochemicals from nuts remains a challenging task and the active constituents include bhilawanols, cardol, anacardic acid, semecarpol and anacardol (Adhami *et al*. 2012; Nair *et al*. 2009). Among these, bhilawanol and anacardic acids are immunogenic, and active exposure causes irritation, blisters, contact dermatitis, and toxicity in humans (Llanchezhian *et al*. 2012).

Due to the high toxicity and challenges in isolating essential bioactive compounds from nuts, we have investigated the other parts of the plant for potential anti-cancer activity. While assessing the plants, we identified leaves that could be used as an alternative source for isolating potential anti-cancer constituents against cancer cells. Hence, it is crucial to focus on how leaves can be used as an alternative and eco-friendly source without altering biodiversity.

In this study, we showed that the ethyl acetate extract of *S. anacardium* Linn. leaves induces apoptosis and cell cycle arrest in the G1 phase, thus inhibiting breast cancer progression. Further analysis revealed that the oral administration of the extract inhibits tumor growth, delays tumor-associated death, and significantly enhances survival in tumor-bearing mice. Therefore our results show the potency of *S. anacardium* leaves extract as a raw material for future anti-cancer drug discovery for breast cancer. Thus, this study established the anti-cancer property of leaves for the first time with less toxicity and efficacy than the nuts.

## 2. Results

Our analysis of plant extract from different parts (root, bark, leaf and fruit) revealed that the leaf is an excellent source for isolating potential phytochemicals that showed higher cytotoxic activity against cancer cells while maintaining normal cells at the baseline level (Fig S1).

### 2.1 Identification of phytochemical constituents from leaf extract

To identify active compounds from *S. anacardium*, we analyzed the ethyl acetate leaf extract in GC-MS and identified seventeen major compounds (Table S1). Among these, (E)-octadec-9-enoic acid had the highest peak area (11.30%). The other potential constituents include, palmitic acid (9.89%), (E)-3,7,11,15-tetramethylhexadec-2-en-1-ol (8.77%), methyl3-(3,5-di-tert-butyl-4-hydroxyphenyl) propanoate (8.06%) and other phytochemicals were listed in the table S1. Compound 1 and 11 belongs to alkaloids, 13, 14, and 15 are steroids, 2 is terpenoids, 15 and 16 are phenolic compound, 17 is a flavonoid, while the remaining are identified as aliphatic compounds.

### 2.2 *Semecarpus anacardium* Linn. leaf extract exhibits anti-cancer activity in cancer cells

The evaluate anti-cancer activity, we treated either petroleum ether (BLPE), ethyl acetate (BLEA) or methanol (BLM) derived leaf extract in six different cell lines, including MCF7, MDA-MB-231, HCT-15, MIN6, EAC and L929. The concentrations of the extracts ranging from 0 to 200 μg/ml were used in our study. Among these, BLEA-derived leaf extract induced the highest cytotoxicity among all cancer cell lines, including MCF-7 and EAC cells, with an IC50 of 0.57 and 1.71 μg/ml, whereas HCT-15, MIN-6 and HepG2 cells showed most minor sensitivity with an IC50 of 6.66, 15.77 and 44.93 μg/ml towards the extracts (Fig 1 and Table S1). Moreover, BLEA showed very low cytotoxicity in normal L929 cells (IC50 value 29.62 μg/ml), indicating that the extracts were less toxic for normal proliferating cells. Therefore, BLEA was selected against MCF-7 cells for further studies (Table S1). Additionally, we compared the cytotoxicity of BLEA with paclitaxel in MCF-7 cells. However, paclitaxel significantly affects the cancer cells at a low concentration compared to BLEA, suggesting that BLEA is a fraction of crude extract (Fig S3). These data indicate that BLEA-derived leaf extract can be a potential alternative source for cancer treatment.

**Figure 1:**
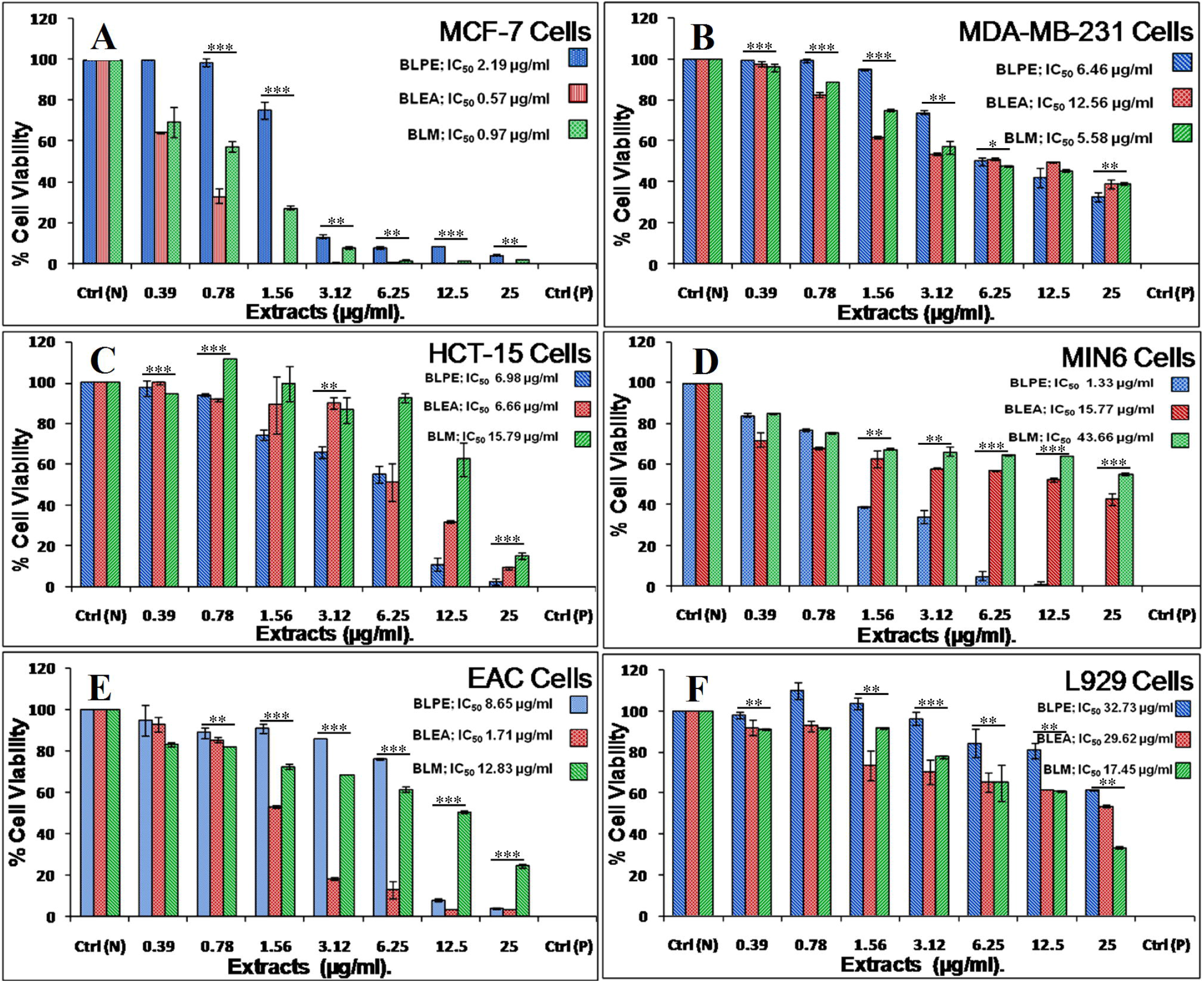
Cytotoxicity assay of *Semecarpus anacardium* leaf extract in different cell lines. (A-F) Histogram showing cytotoxic assay of leaf extract in (A) Human breast cancer cells (MCF-7), (B) Human triple-negative breast cancer cells (MDA-MB-231), (C) Human colorectal carcinoma cells (HCT-15), (D) Mouse insulinoma cells (MIN6), (E) Mouse ascetic carcinoma cells (EAC) and (F) Mouse fibroblast cells (L929) treated with BLPE, BLEA & BLM and cytotoxicity was evaluated by MTT assay after 48 hours of treatment. BLEA-derived leaf extract showed a robust response of cell viability in MCF-7 cell lines compared to others. The graphs were plotted using the mean of the data and error bar depicting □±□SEM (*p*-value: *≤0.05, **≤0.025, ***≤0.01).

### 2.3 *Semecarpus anacardium* Linn. leaf extract alters cell morphology and induces apoptosis in cancer cells

Establishing proper cell morphology is an essential characteristic feature of healthy cells. To assess cellular morphology, we treated MCF-7 cells with BLEA and examined them under a light microscope to observe if BLEA affects any morphological changes in cancer cells. PTX-treated sample was taken as a positive control to compare cellular morphology with BLEA. Our microscopic analysis revealed cytoplasmic vacuolization in the BLEA and PTX-treated cells, while untreated control did not show such structures (Fig. 2 A-C). An earlier report has indicated that the cells undergoing stress accumulate vacuolar structures in the cytoplasm. The accumulation of vacuoles and changes in cellular morphology might induce a pathway leading to cell death. Hence, we speculate that apoptosis might be induced in BELA-derived leaf extract. To evaluate the cell death, MCF-7 cells were grown in coverslips, stained with DAPI and examined under a fluorescence microscope. Our microscopic analysis revealed a significant alteration in the nuclear morphology, including chromatin condensation, nuclear fragmentation and shrunken nuclear morphology (Fig 2 D-F).

**Figure 2:**
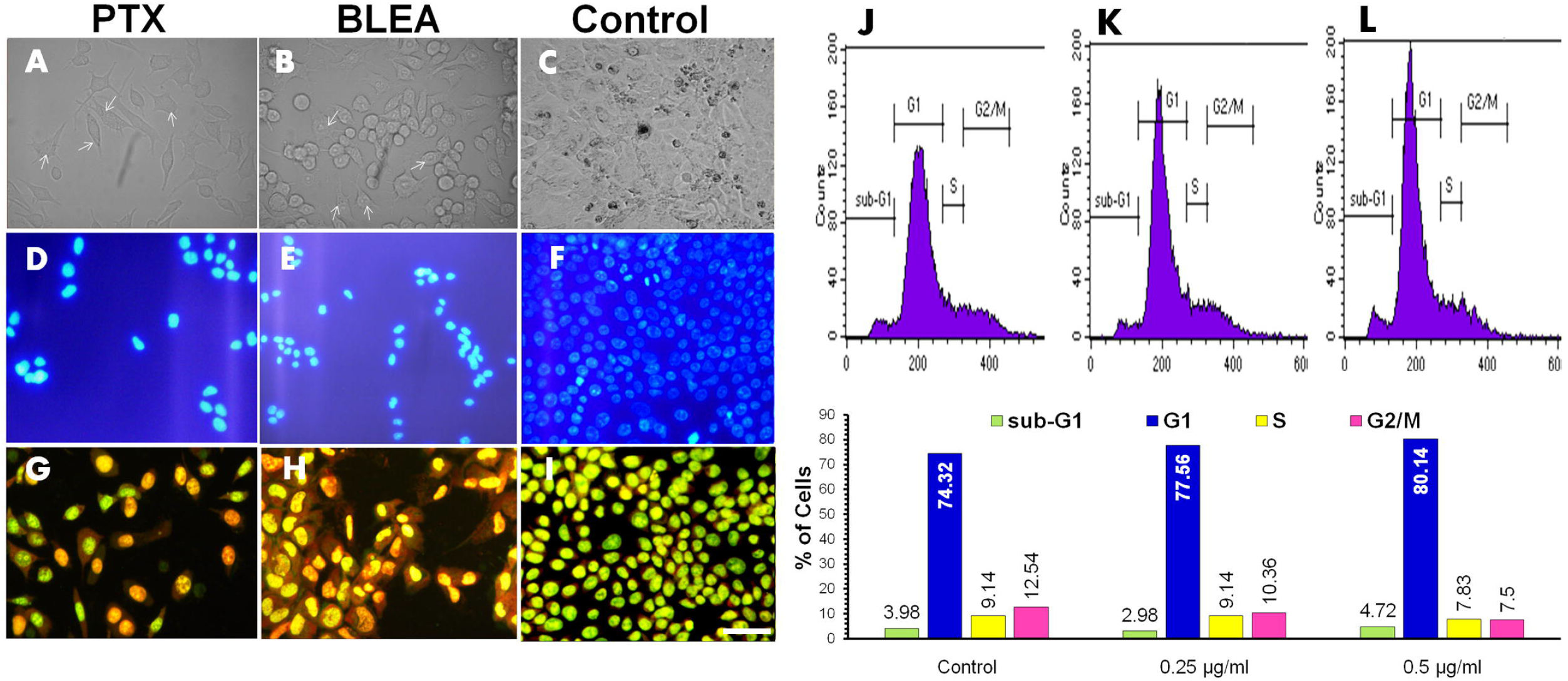
BLEA affects cell morphology, apoptosis and cell cycle arrest in MCF-7 cells. (A-C) Representative light microscopic image showing vacuolization in cells treated with (A) Paclitaxel (2.13 ng/ml), (B) BLEA (0.57 μg/ml) and (C) control. (D-F) Representative fluorescence microscopic image showing the evaluation of nuclear morphology in cells treated with (D) Paclitaxel (2.13 ng/ml), (E) BLEA (0.57 μg/ml) and (F) control cells. (G-I) The cell death was analyzed by double staining with ethidium bromide and acridine orange in cells treated with (G) Paclitaxel (2.13 ng/ml), (H) BLEA (0.57 μg/ml) and (I) control cells. (J-L) The BLEA-treated cells were further analyzed in FACS and showed cell cycle arrest in the G1 phase in different concentrations of leaf extract in (J) control cells, (K) 0.10 μg/ml and (L) 0.20 μg/ml. The scale bar represents 20 μm.

Moreover, our ethidium bromide and acridine orange staining data indicate that BLEA potentiates apoptosis by inducing nuclear fragmentation, nuclear contraction, and cytoplasmic membrane blebbing in cancer cells (Fig 2 G-I). The entry of ethidium bromide into the cells leads to a change in orange to red color fluorescence, indicating that cells are undergoing apoptosis. Together, these data indicate that BLEA mediates apoptosis by inducing stress in cancer cells.

### 2.4 BLEA-derived leaf extract induces cell cycle arrest in cancer cells

Nevertheless, cancer cell exhibits induction of S and G2/M phase transition leading to cancer progression. Moreover, we analyzed the cell cycle progression from G1-S to G2/M transition to examine if BELA-derived leaf extract affects cell cycle progression. Our analysis revealed that BLEA-derived leaf extract modestly increased the population of cells at the G1 phase. Furthermore, the sub-G1 population of cells was also modestly increased at the higher concentration of BLEA (500 ng/ml). Conversely, the percentage of cells in the S and G2/M phases decreased with an increase in BLEA concentration (Fig 2 J-L). These observations suggest that BLEA induces cell cycle arrest at the sub-G1 and G1 phases.

### 2.5 *Semecarpus anacardium* Linn. leaf extract inhibits cancer cell migration

Cellular migration in cancer cells is essential, enabling them to metastasize. Wound healing has been widely used as a model to examine cancer cell migration. Hence, we performed a wound healing to assess the effect of BLEA on cancer cell migration. The MCF-7 cells were treated with BLEA and examined on the day 1st, 3rd and 5th for wound healing. Interestingly, we observed a significant decrease in cellular migration, resulting in delayed wound healing compared to control cells (Fig. 3). This data suggests that BLEA-derived leaf extract significantly reduces migration and progression in cancer cells.

**Figure 3:**
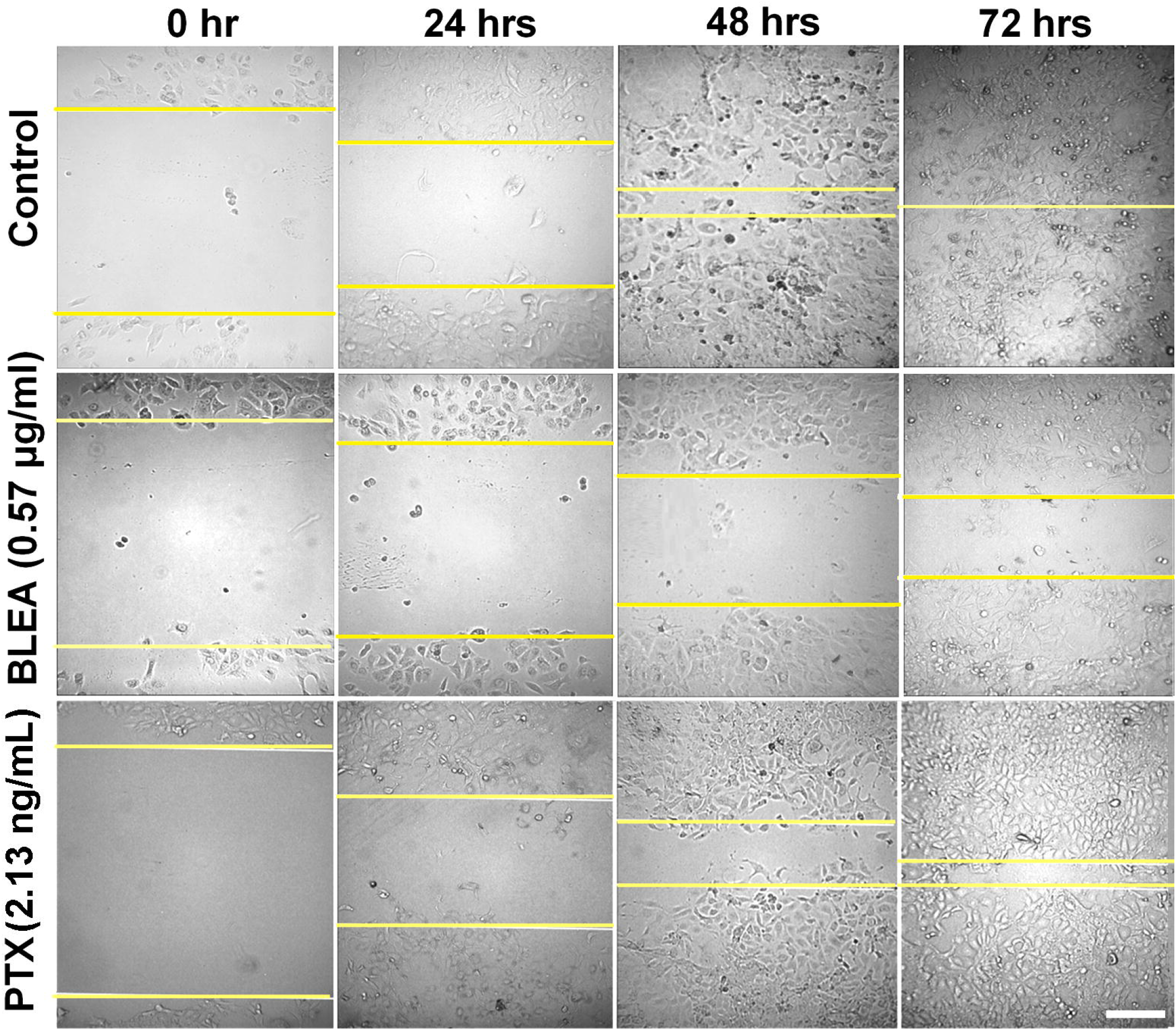
BLEA inhibits cellular migration in MCF-7 cells. The wound healing assay was used to examine the effect of BLEA on wound closure and cellular migration. The images were captured in the light microscope with 40X magnification. Note that BLEA-derived leaf extract significantly suppressed cancer cell migration compared to PTX-treated and control cells. The scale bar represents 10 μm.

### 2.6 BLEA-derived *Semecarpus anacardium* Linn. leaf extract does not show any toxicity in mice

Our in vitro cell culture experiments could not differentiate if BLEA-derived leaf extract has any consequence on the toxicity of animals. Hence, to gain insights into it, we evaluated the toxicity by oral administration of BLEA in mice using the standard protocol (test number 425 of OECD). Interestingly, BLEA did not show any toxicity in mice, even at high doses. Next, we analyzed the tissues of vital organs to see any alteration of cellular architecture due to toxicity. However, careful examination of the histological section revealed any notable alteration in cellular architecture. Besides, we did not see any body weight loss, indicating that BLEA is non-toxic to the animals for oral administration (Fig. S4 and Table S2).

### 2.7 BLEA inhibits tumor progression in mice

Next, we examined the anti-cancer potency of BLEA using EAC (Ehrlich ascites carcinoma) cells induced tumor model in mice. Based on a pilot study, a 100 mg/kg body weight dose of BLEA was selected to evaluate anti-tumor activity in mice (Fig S5). The treatment of tumor-bearing mice started from the 8th day of EAC cell injection, when tumors became measurable, with a daily dose of 100 □ mg/kg body weight of BLEA till the 28th day. Next, we monitored the tumor progression on alternative days. Interestingly, the oral administration of BLEA resulted in a significant decrease in tumor volume (58.67, 224.00 and 298.67□mm^3^) compared to the control groups (Fig. 4B-D). In contrast, the mice that did not receive any treatment for BLEA, the mean tumor volume was significantly increased (58.67, 388.98 and 780.44□mm^3^) on the day 8th, 18th and 28th compared to treated groups (Fig. 4A). Additionally, the histological sections of thigh, liver and spleen from tumor-bearing mice revealed alterations in cellular morphology compared to the treated and control mice (Fig. 4F-H). Altogether, these data suggest that BLEA has the potency to reduce tumor progression significantly in mice. Therefore, BLEA can be used in vivo and in vitro models to study the nature of cancer progression.

**Figure 4:**
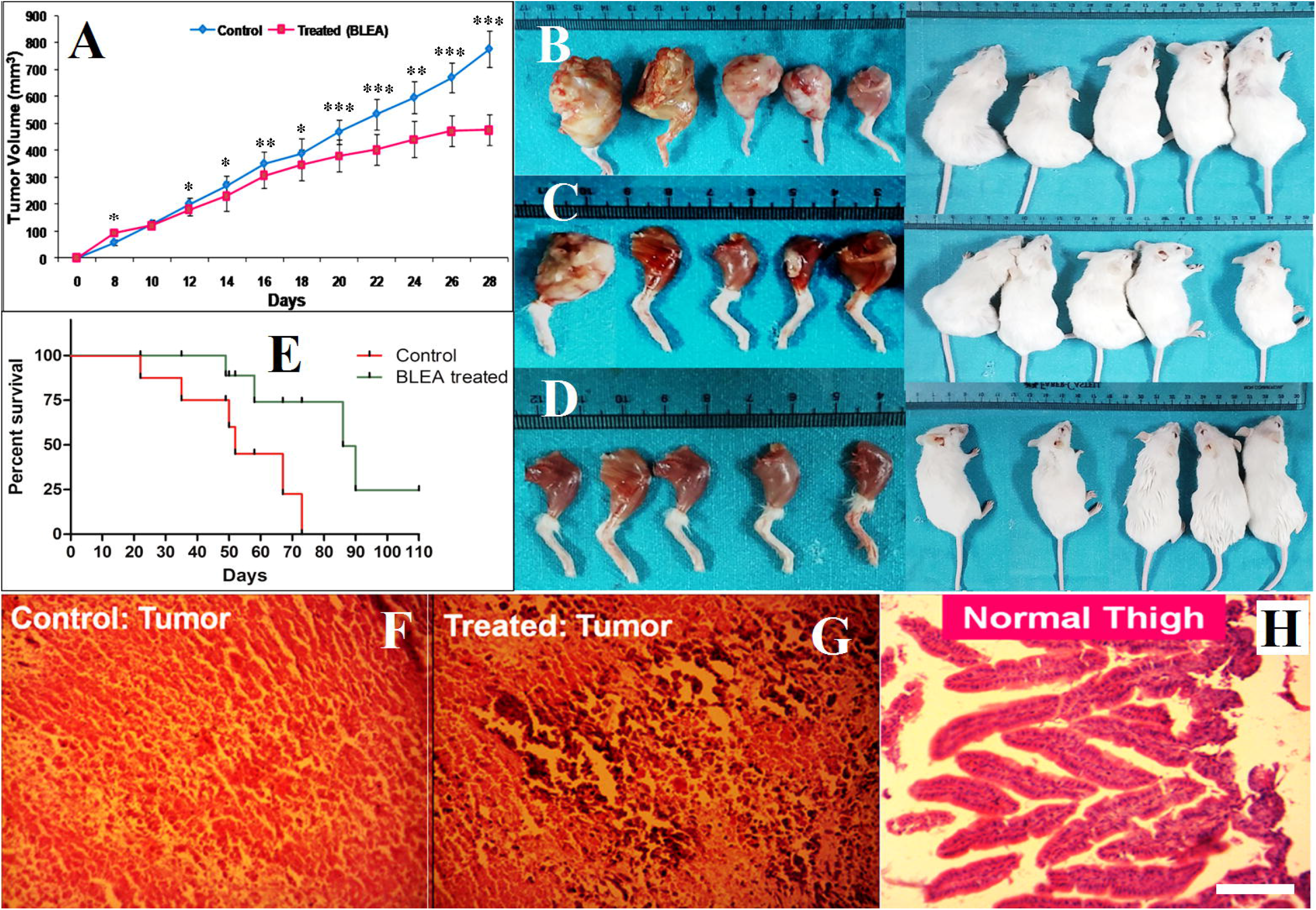
BLEA inhibits tumor progression and enhances longevity in tumor-bearing mice model. The EAC cells were injected in the left thigh to develop a solid tumor in mice and 100 mg/kg body weight dose of BLEA was administered orally after eight days of EAC injection throughout the experimental period. Graphical analysis showing tumor volume in (A) BLEA treated and untreated tumor-bearing mouse model. The graphs were plotted using the mean of the data and error bar depicting □±□SEM (*p*-value: *≤0.05, **≤0.025, ***≤0.01). (B-D) Representative light microscopic image showing the size of tumors in (B) control tumor-bearing group, (C) BLEA treated tumor group and (D) thigh of normal mice. Note that oral administration of BLEA significantly restored tumor volume to the thighs of normal mice. (E) Longevity curves (Kaplan–Meier) of treated and untreated tumor-bearing mice showed significant enhancement in survival rate in BLEA-treated tumor-bearing mice. (F-H) Represented histological section showing (F) untreated HE-stained tumor, (G) section of treated HE-stained tumor and (H) normal thigh section. Note that BLEA-treated tumor shows a significant number of cell deaths compared to control tumors. (F-G) The scale bar represents 20 μm.

### 2.8. BLEA delays tumor-associated death and enhance the survival of tumor-bearing mice

Cancer progression significantly reduces the life span of humans. Hence, we speculate that BLEA-derived leaf extract might potentially improve the life span of animals. Notably, our data showed a significant delay in tumor-associated death in the mouse. Moreover, the first death in BLEA-treated mice was recorded on day 49th. In contrast, the first death in control tumor-bearing mice was recorded on the 22nd.

Additional studies revealed that BLEA induces an increment in survival rate (~50%) in tumor-bearing mice compared to untreated tumor-bearing animals. The untreated tumor-bearing control survived only up to 73 days after EAC injection, whereas the BLEA-treated tumor-bearing mice survived more than 110 days. Besides, further study revealed a significant decrease in the number of EAC cells in tumor-bearing mice compared to untreated control (Fig. 4E). Therefore, our study identified the most efficient method to isolate and evaluate the anti-cancer properties of BLEA in vivo model. Overall, our results showed that BLEA exhibits cytotoxicity by inducing apoptosis, including enhancing the survival rate of tumor-bearing mice. Thus, this study suggests that BLEA could be a prime source for developing a chemotherapeutic agent to prevent cancer progression in humans.

## 3. Discussion

The development of chemoresistance among cancer cells is a major concern in the treatment, leading to a high mortality rate in cancer patients. The inactivation, alteration and efflux of drug targets influence the chemoresistance of cancer (Arya *et al*. 2015). Therefore, plant extract or a mixture of multi-targeted compounds with anti-tumor activity are potent tools to overcome drug resistance. Traditionally used medicinal plants serve as the prime source for developing chemotherapeutic drugs with minimal side effects. Hence, it reduces the time and cost of developing potential chemotherapeutic agents.

*Semecarpus anacardium* is a widely studied medicinal plant, which has been described in Ayurveda for the preparation of several poly-herbal formulations to treat clinical ailments. For instance, inflammation, neurological disorders, problems associated with geriatric, vitiligo, cardiac troubles, spleen enlargement, hemorrhoids, rheumatism, diabetes, baldness, etc., including cancer (Arya *et al*. 2015). The nut extract regulates the activity of a critical enzyme involved in carbohydrate metabolism, which is elevated in cancer cells during tumor growth to enhance the synthesis of nucleic acid and lipids (Semalty *et al*. 2010; Sharma *et al*. 1995). Besides, *S. anacardium* extract has been studied in hypoxia-associated factors and extracellular matrix-associated enzymes in tumors, inhibiting tumor progression and metastasis in DMBA-induced tumor-bearing rats (Sharma *et al*. 1995) (Arya *et al*. 2015). Additionally, the milk of nuts has been reported to have an anti-cancer property in hepatocellular carcinoma in Wistar rats (Joseph *et al*. 2013). However, the nut extract is cytotoxic and induces cell death through apoptosis in breast cancer (T47D) cells by arresting the transition of the G2/M phase (Mathivadhani *et al*. 2007a). The compounds isolated from nuts show cytotoxicity with an IC50 value range from 0.10 – 5.9 μg/ml among various cancer cell lines. Besides, 3-O-methyl quercetin and kaempferol show cytoprotective properties against free radical-induced cellular damage. In contrast, the isolated compound shows a synergistic effect with doxorubicin and induces apoptosis via arresting cells in the G2/M phase of the cell cycle in cancer cells (Kumar *et al*. 2016; Nair *et al*. 2009).

In this study, we evaluated the anti-cancer properties of *S. anacardium* as an alternative source of phytochemicals for therapeutic use. However, the medicinal application of leaves remains largely unknown. Moreover, the leaf extracts were prepared and evaluated for anti-cancer activity using various cell lines and animal tumor models. The ethyl acetate extract showed the most potent cytotoxic activity against different cancer cell lines and induced apoptosis in MCF-7 cells. However, it did not show any alteration of activities in normal proliferating L929 cells. Besides, BLEA exhibited cell cycle arrest in the G1 phase and apoptotic cell death in MCF-7 cells.

Additionally, the oral administration of BLEA induces significant tumor growth suppression and enhances the longevity of tumor-bearing mice. The IC50 values of the leaf extracts were determined and found to be lower than the compounds isolated from nuts and considered to be more toxic than leaf extract studies (Nair *et al*. 2009). For instance, the IC50 values of leaf extract derived in petroleum ether were 1.33-8.65 μg/ml, while ethyl acetate and methanol-derived leaf extract exhibited 0.57 – 15.77 μg/ml and 0.97 – 43.66 μg/ml, respectively. In contrast, the nuts extracts exhibited higher IC50 values and were found to be 32.73, 29.62 and 17.45 μg/ml when extracted in petroleum ether, ethyl acetate and methanol, respectively. Besides, the previous report has indicated that a selectivity index is an essential tool in bioassay-guided fractions. Hence, we sought to calculate the selectivity index, which ranges from 0.40 to 52.17 for all three extracts in different cancer cells. Among these, the selectivity index of ethyl acetate leaf extract is higher in human breast carcinoma (MCF-7) cells. Hence we selected BLEA and MCF-7 cells for further studies.

Nevertheless, chemotherapeutic agents can affect cell cycle progression and promote apoptosis-mediated cell death (Hanahan and Weinberg 2011). Several chemotherapeutic drugs have been tested that promote apoptosis and inhibit tumor progression and are considered novel anti-cancer drugs (Singh *et al*. 2016). Our study revealed the formation of many cytoplasmic vacuoles following treatment with BLEA in breast cancer (MCF-7) cells, indicating the induction of apoptosis-like cell death (Mallick *et al*. 2015; Vijayarathna *et al*. 2017). Double staining with acridine orange and ethidium bromide labels untreated cells, early apoptotic cells, late apoptotic cells and necrotic cells, indicating BLEA-induced apoptosis in MCF-7 cells. Acridine orange is permeable to the intact plasma membrane and can bind DNA, resulting in green fluorescence. In contrast, Ethidium bromide enters the cells only with a broken plasma membrane, binds to nucleic acid and emits red fluorescence. The green, yellow, orange and red fluorescence represent normal, early apoptotic, apoptotic and late apoptotic/necrotic mediated cell death (Ramya *et al*. 2018). Besides, the BLEA-treated cells showed significant changes in cell morphology, which further confirmed the induction of the apoptosis mode of cell death (Ramya *et al*. 2018). Moreover, we observed that BLEA alters the cell cycle and arrests the MCF-7 cells at G1 phase. Interestingly, the nut extract has been reported to arrest cancer cells at the G2/M phase, suggesting that nut and leaf extract differ in their mode of action in cancer cells (Krishnan *et al*. 2017; Mathivadhani *et al*. 2007a). A wound healing assay is considered an essential tool for studying cell migration and a model for cancer metastasis. Numerous phytochemicals and herbal extracts have been reported to inhibit cellular migration, including preventing metastasis in cancer patients (Gao *et al*. 2018; Jiang *et al*. 2017; Liang *et al*. 2015). Our study reported that BLEA-derived leaf extract inhibits cell migration, indicating the anti-cancer properties of BLEA.

The different extracts of *Semecarpus anacardium* nuts are thoroughly studied for their toxicity in humans and other animal models (Llanchezhian *et al*. 2012; Patwardhan *et al*. 1988; Sundaram *et al*. 2018). However, the purification process remains more challenging to reduce toxicity before clinical use (Maurya *et al*. 2015). In contrast, this study determined that ethyl acetate leaf extract showed no toxicity during oral administration in mice. Besides, we did not observe any physiological or behavioral changes, even in mice receiving a high dose (5000 mg/kg b. wt.) of the BLEA. Furthermore, histological analysis of stomach, liver, kidney and lung tissues revealed no changes in cellular structure in BLEA-treated mice.

Moreover, our study has further extended to evaluate in vivo anti-tumor activity in the EAC cells-induced tumor model. The EAC cells are mouse-derived ascetic carcinoma cells readily accepted in mice bodies with high transplantable efficacy, rapid growth and confirmed malignancy. The subcutaneous injection of EAC cells resulted in the development of a solid tumor in mice (Ezzat *et al*. 2018; Mishra *et al*. 2018b). In this report, we evaluated the anti-tumor potency of BLEA in EAC-induced allograft tumors, which showed significant inhibition of tumor growth and increased the survival of tumor-bearing mice. These results are consistent with the other herbal extracts, for instance, *Achras sapota, Vernonia condensate, Eclipta alba*, etc. (Arya *et al*. 2015; Srivastava *et al*. 2014; Thomas *et al*. 2016). Moreover, the histological sections of the tumor showed apparent death of tumor cells induced by BLEA (Srivastava *et al*. 2014). Therefore our study explored a novel method to isolate and evaluate potential phytochemicals from the leaf with minimal toxicity for normal cells while exhibiting anti-cancer activity in cancer cells.

Moreover, BLEA inhibited tumor growth, delayed death and significantly enhanced survival in the tumor-bearing mice model. Therefore, BLEA is a potential extract for the development of anti-cancer chemotherapy. However, additional studies are necessary to develop and elucidate molecular pathways involved in developing an efficient anti-cancer drug for *Semecarpus anacardium*.

## 4. Materials and Methods

### 4.1 Collection of plant specimens and phytochemical analysis

The leaves of *Semecarpus anacardium* Linn. were collected from Banaras Hindu University, Varanasi, India and identified by the Department of Dravyaguna (Ayurvedic Pharmacognosy & Pharmacology), Faculty of Ayurveda, Institute of Medical Sciences, Banaras Hindu University, Varanasi, India (Accession Number: DG/18/171). The plant materials were dried at room temperature for two weeks and minced into a coarse powder through the domestic electrical grinder. Three times, one kilogram of leaves powder was macerated up to 24 hours with continuous stirring in 5 liters of petroleum ether, ethyl acetate and methanol. The supernatant was filtered through Whatman filter paper of 11μm pore size. Further, each filtrate was dried by a rotary vacuum evaporator at reduced pressure (Singh *et al*. 2020). The ethyl acetate extract of *S. anacardium* leaves (SLE) was assessed by Gas chromatography-mass spectrometry (GC-MS), the JEOL GCMATE II GC-MS at SAIF, IIT Madras using standard procedure (Usharani and Vasudevan 2017). The data were matched with the standards (Elumalai *et al*. 2015).

### 4.2 Cell culture

Following cell lines such as human breast adenocarcinoma (MCF-7) cells, human triple-negative breast carcinoma (MDA-MB-231) cells, human colon adenocarcinoma (HCT-15) cells, mouse insulinoma (MIN-6) cells, mice undifferentiated breast carcinoma (Ehrlich-Lettre ascites carcinoma, EAC) cells and mice fibroblast (NCTC clone 929, L929) cells were procured from National Centre for Cell Sciences (NCCS), Pune, India. The cells were cultured in DMEM with high-glucose, sodium bicarbonate, L-glutamine and sodium pyruvate supplemented with 10% fetal bovine serum (FBS) and 1% antibiotics-antimycotic (10000 unit/mL of penicillin G, 10 mg/mL of streptomycin sulfate and 25 μg/mL amphotericin B in 0.9% normal saline) solution at 37 °C in a humidified incubator with 5% CO2. About 70-80% of confluent cells at the log phase were harvested by trypsinization (1X Trypsin-EDTA Solution, Himedia) (Singh *et al*. 2020). MCF-7 cells were used throughout the study, while EAC and L929 cells were used in the cytotoxicity assay.

### 4.3 Cytotoxicity assay

For the MTT assay, cells were seeded in a 96-well culture plate at 1 X 10^4^ cells/ml density and incubated overnight. The culture media was replaced with fresh media containing leaf extracts and incubated for 48 hours. After 48 hours, 100 μl of 3-[4,5-dimethylthiazol-2-yl]-2,5-diphenyl tetrazolium bromide (MTT) solution at a concentration of 0.5 mg/ml was added to the cells and kept it for 4 hours at 37 °C. The treated cells were centrifuged at 750 rpm for 5 minutes (Remi R8C) and the supernatant was replaced with 100 μl DMSO to solubilize the formazan crystal. Next, we measured the absorbance at 570 nm in a microtitre plate reader (Biorad, India) (Mishra *et al*. 2018a; Sekhar *et al*. 2019; Singh *et al*. 2020). The cell viability was calculated as the percentage of cell viability = (OD of the treated cells /OD of the untreated cells as control) × 100. Further, IC50 was calculated using standard procedures.

## 5. Authors contribution

RKS and AKS designed the research; RKS, AR, RT, SSV, VS and SCG performed the research; RKS and AKS analyzed the data; BM and RKS wrote the paper.

## 6. Acknowledgment

We thank Banaras Hindu University, Varanasi, India, for its infrastructure facilities. RKS thanks BHU and UGC for fellowships.

## 7. Declaration

All the authors are responsible for the part of the data represented in the paper, as mentioned in the authors contribution section. The authors declare no compiting financial interest

## Notes

### Competing Interest Statement

The authors have declared no competing interest.

### Summary of Updates

Dear BIORXIV Staff, There are significant revisions have been made throughout the manuscript in the following part: 1. Introduction 2. Results 3. Discussion A significant part of the materials and sections has been moved to the supplementary materials. These include rephrasing words and sentences. Figure 5 has been removed from the manuscript. Figure S2 has been removed from the supplemental file. Bhagaban Mallik has been added as a new author in the current version.

