## Supplemental Materials for "*Semecarpus anacardium* Linn. leaf extract exhibits activities against breast cancer and prolongs the survival of tumor-bearing mice"

**Abstract**

*Semecarpus anacardium* Linn. is commonly used in various traditional medicines from ancient times. The nuts have been described in Ayurveda medication systems to treat numerous clinical ailments. However, isolating phytochemical constituents from nuts remains challenging and exhibits cytotoxic effects on other cells. In this study, we have standardized procedures for isolating phytochemicals from the leaf extract. The ethyl acetate leaf extract selectively affects cancer cells in a dose-dependent manner (IC50: 0.57 µg/ml in MCF-7 cells) in various cancer cell lines.

Next, we examined if the extract incubation could induce cell cycle arrest and suppress cell migration in the cell culture model. Consistent with this idea, the leaf extract could potentially affect the aggressive migration nature of cancer cells. Moreover, oral administration of extract significantly restored tumor growth in mice. Together, these observations suggest the anti-cancer activities of *S. anacardium* leaf potential for both in vitro and in vivo models.

Keywords: *Semecarpus anacardium*; Ayurveda; Breast cancer; Apoptosis; Cell migration; Phytochemical.

**Experimental Methods**

**4.1 Analysis of cell morphology**

For microscopic analysis, MCF-7 cells were seeded into 96 wells of culture plates. Next, cells were treated with the leaf extract for 24 hours and observed under the inverted light microscope (Dewinter, India) with a 40X objective lens (Singh *et al.* 2018; Singh *et al.* 2020). Further, acridine orange (AO) and ethidium bromide (EtBr) was used to detect cell death mode. Acridine orange is a cell-permeable dye that binds with the nucleic acid to emit green fluorescence, while ethidium bromide can enter the cell membrane and binds to the nucleic acid to emit red fluorescence. Paclitaxel was used as a positive control, while 10 mM PBS was taken as a negative control. Next, we removed the culture media and the cells were washed with 10 mM PBS (pH 7.4) and incubated with 100 μg/ml AO and 100 μg/ml EtBr in the dark for 30 minutes. Finally, the cells were washed with 10 mM PBS (pH 7.4) and examined under the inverted fluorescence microscope (Dewinter, India) using a 40× objective lens (Singh *et al.* 2018)

**4.2 Analysis of nuclear morphology**

Human breast cancer (MCF-7) cells were seeded at 1 X 104 cell density and incubated for 24 hours at 37 °C in a humidified incubator with 5% CO2. The seeded cells were treated with leaf extract for 48 hours and stained with DAPI (4′-6-diamidino-2-phenylindole) (Genetix, India) to analyze nuclear morphology (Singh *et al.* 2018). The cells were washed with 10 mM PBS (pH 7.4) and mounted on a slide for microscopic analysis. The images were captured in an inverted fluorescence microscope (Singh *et al.* 2020).

**4.3 Cell cycle analysis**

Briefly, the cells were incubated with leaf extracts and washed with 1X PBS before cell cycle analysis. Further, the cells were stained with 70% chilled methanol containing RNase A and propidium iodide. Next, FACS was performed in Cell Quest software (Becton Dickinson) to determine the population of cells in different phases of the cell cycle.

**4.4 Wound healing assay**

MCF-7 cells were grown in DMEM culture up to 80% confluency, trypsinized and seeded into a 6-well plate (1 × 10^5^ cells/ml) at 37 ˚C for 24 hours (Mishra *et al.* 2018a; Sekhar *et al.* 2019). The culture media were replaced with fetal bovine serum-free media for 24 hours and a wound was created with a sterile 20 µl pipette tip following washing with 1X PBS. Further, the wound-containing cells were treated with leaf extract in a 1 ml complete medium in the test and control groups. The cells were examined at 24, 48 and 72 hours intervals. Images were captured and analyzed for cellular migration in an inverted microscope (Dewinter, India) equipped with a 10X objective lens (Singh *et al.* 2020).

**4.5 Animal model and ethical statement**

For in vivo studies, Swiss albino mice of either sex weighing 18–22 grams of about 8–10 weeks old were procured from the central animal facility, Institute of Medical Sciences, Banaras Hindu University, Varanasi, India, with approval of the institutional ethical committee (Dean/2016/CAEC/337). The animals were kept in polypropylene cages and provided water and a standard pellet diet (Agro Corporation Pvt. Ltd., Bengaluru, India). The mice were accommodated under controlled temperature (27 ± 2 ˚C) and humidity with a 12 hours light and dark cycle by following CPCSEA guidelines for the experimental study.

**4.6 Anti-tumor evaluation**

To determine anti-tumor activity, 18 tumor-bearing mice and six normal mice were divided into four groups. Mice of group I, group II and group III were administered 20 mg/kg body weight Paclitaxel, 100 mg/kg bodyweight BLEA (suspended in water) and an equal volume of water. In contrast, group IV was normal mice without tumors. All mice of group I and Group II were given respective drugs regularly throughout the experimental period for 28 days. The control mice of group IV were given any treatment for 28 days. Tumor size was measured using vernier calipers on alternative days for tumor animals and tumor volume was calculated using the formula V = 0.5 × a × b2, where 'a' and 'b' indicates the major and minor diameter, respectively. At the end of the 28th day of the experimental period, animals from each group were sacrificed to collect kidneys, liver, thigh and spleen from Paclitaxel, BLEA-treated tumor-bearing and normal mice. The morphological changes were evaluated using hematoxylin-eosin-stained histological slides (Srivastava *et al.* 2014).

**4.7 Animal survival study**

For longevity assay, the effect of BLEA on the life span of the tumor-bearing mice was assessed. In this study, 12 mice received EAC cells in their left thigh to induce the tumor. They were further divided into two groups (6 mice in each). Group I served as tumor control and group II received oral administration of BLEA 100 mg/kg body weight daily after seven days of tumor (EAC) cell injection till the survival of the last animal. The life span of BLEA-treated mice was calculated and compared with untreated tumor-bearing animals. The life span of experimental animals was monitored and calculated by using the following formula [(T − C)/C] × 100, where 'T' indicates the number of days the treated animals survived and 'C' indicates the number of days that tumor animals survived (Arya *et al.* 2015).

**4.8 Histology**

The tissues were collected from control and BLEA-treated animals and processed for histological evaluations (Thomas *et al.* 2016). The excised tissues were fixed in 4% neutral saline formalin, dehydrated in alcohol and embedded in paraffin wax. The thin section (5 μm) was made in Microtome (Leica Biosystems, Germany) and then stained with hematoxylin and eosin. Images of all tissue sections were captured in a light microscope (Dewinter).

**4.9 Statistical analysis**

The data were analyzed using GraphPad Prism 6.0 as described previously (Mallik *et al.* 2017; Raut *et al.* 2017) The analysis of variance was performed on three independent cytotoxicity results by one-way ANOVA using the post hoc Tukey test. The results were considered significant at a *p*-value <0.05. The data were represented as mean±SD.

**4.10 Evaluation of the acute oral toxicity**

The evaluation of acute oral toxicity was carried out per the procedure described by the Organization for Economic Cooperation and Development (OECD test 425). A total of 18 mice of either sex were divided into three groups (3 male & 3 female in each group) randomly, such as experimental groups 1 & 2 and control group 3, respectively. A day before an experiment, animals were kept on fasting. The next day, the ethyl acetate extract was administered orally using a 16G oral round ball tip feeding curved needles to the mice of the experimental groups. A single dose of 2000 mg/kg and 5000 mg/kg body weight were administered among the mice of experimental groups 1 and 2, respectively, while distilled water was given to the control group mice. All mice were observed individually after the first 30 min dosing; particular attention was given during the first 4 hours and followed up from 24 hours to the 14th day of the dosing to collect signs and symptoms of toxicity(Amelo *et al.* 2014; Sreeja *et al.* 2018). Careful observation was made to investigate the potential occurrence of tremors, convulsions, salivation, diarrhea, lethargy and drowsiness. The live weight of the animals was monitored on days 0, 7 and 14 of the dosing as one of the toxicity parameters. At the end of the experiment, the animals were sacrificed by cervical dislocation of anesthetized animals. The brain, stomach, liver, spleen, lungs, kidneys, muscles, esophagus and small intestine were collected to evaluate histopathological changes.

**4.11 Tumor model development**

EAC cell-induced tumor model was chosen for this study because of mouse origin, which can be easily transplanted to the immunocompetent mouse. The EAC cells were harvested from a 75 cm^2^ cell culture flask with 70-80% cell confluence by trypsinization and centrifugation at 750 rpm (Remi R8C) for 5 min. The harvested cells were diluted in 10 mM phosphate buffered saline (1X PBS) at a density of 5 × 10^6^ cells, and 200 µl of viable cell solution were injected into the peritoneal cavity of each recipient mouse of 21±1 g body weight and allowed to multiply. The EAC cells were isolated from the peritoneal cavity of the donor mice after 8–10 days of inoculation. Next, the EAC cells were diluted in 1X PBS and centrifuged at 750 rpm (Remi R8C) for 5 minutes. Furthermore, the cell pellet was resuspended in sterile 10 mM PBS and injected (100 µl at 1 × 10^7^ cells/ml) in the left thigh of female mice to develop a solid tumor (Bhattacharjee *et al.* 2017; Mishra *et al.* 2018b).

**4.12 Pilot study for effective dose determination**

After four days of EAC inoculation, a pilot study was set up to determine the effective dose in EAC-induced tumor-bearing mice and divided into four groups of 3 mice each. The tumor-bearing mice were administered 200, 100 and 50 mg/kg body weight of ethyl extract to test the drug's effectiveness in groups 1, 2 and 3. At the same time, 100 µl distilled water was given to the control mice of group 4 for 14 days, along with a regular diet and water ad libitum [53]. At the end of the 14th day, the mice were sacrificed and tumors were dissected to measure tumor volume. Tumor size was measured by Vernier calipers using the formula V = 0.5 × a × b2, where 'a' and 'b' indicates the major and minor diameter, respectively.

**Supplemental Figures**

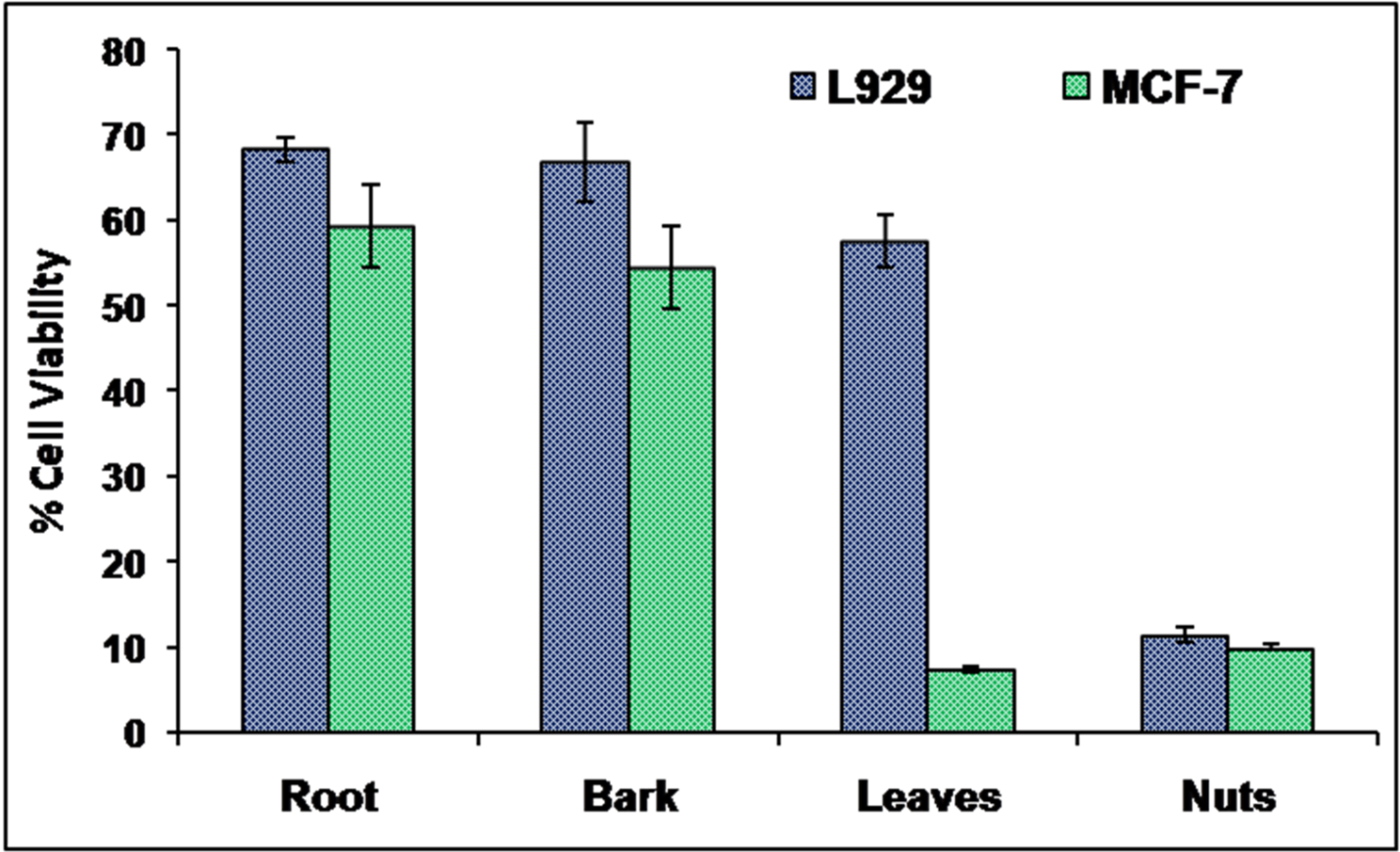

**Figure S1: Cytotoxicity assay of the extract derived from different parts of *Semecarpus anacardium***

The histogram shows the cytotoxicity assay of the extract derived from root, bark, leaves and fruits in MCF-7 and L929 cells. Note that leaf extract showed maximum cytotoxicity in MCF-7 cells compared to the control L929 cells.

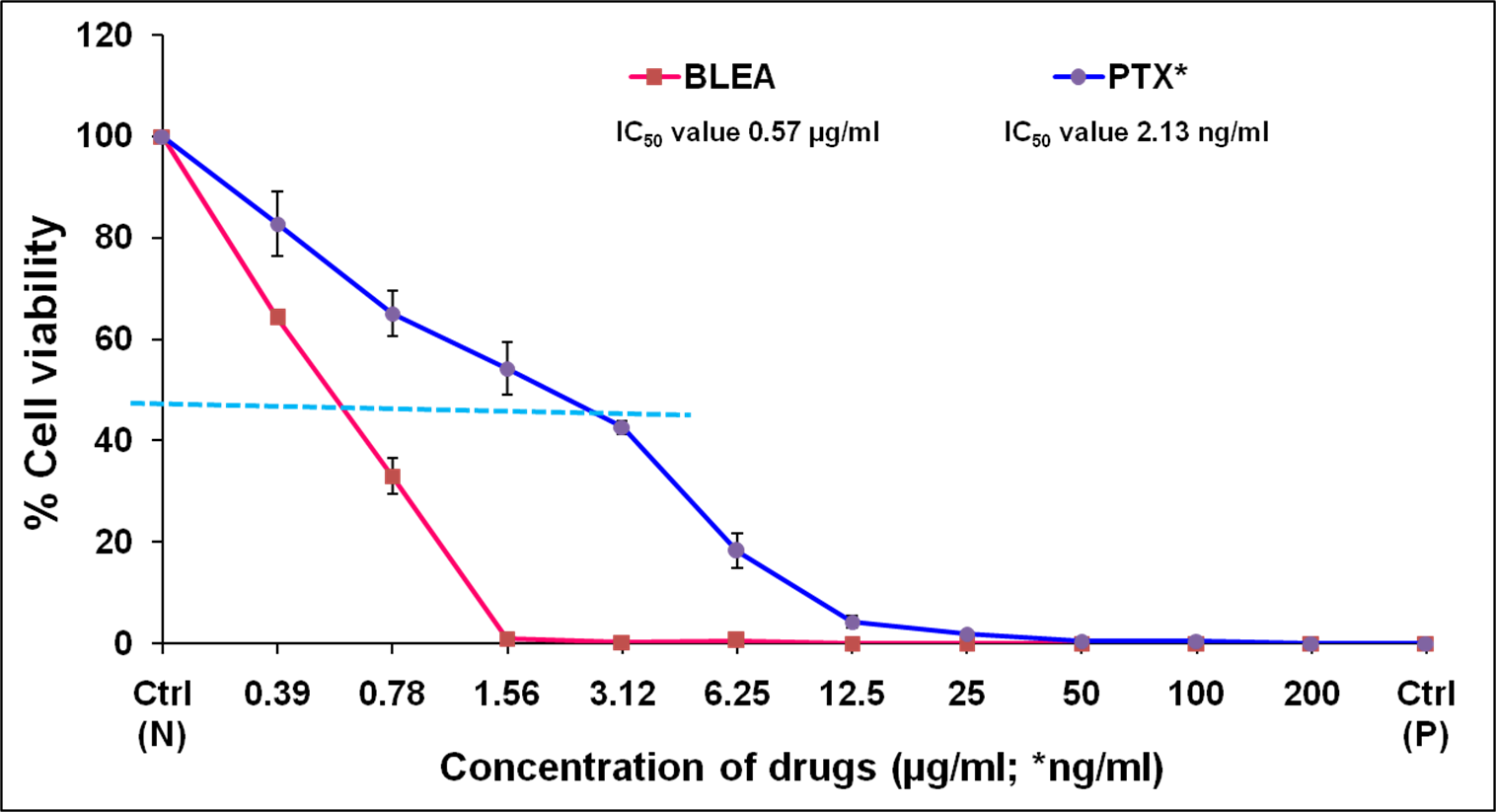

**Figure S2: The BLEA-derived leaf extract shows a lower IC50 value**

The graph compares the cytotoxicity of PTX and BLEA in MCF-7 cells. The IC50 value of BLEA was found to be 0.57µg/ml, whereas Paclitaxel (PTX) was represented as 2.13 ng/ml. Note that BLEA-derived leaf extract exhibits more significant cytotoxicity with a lower IC50 value.

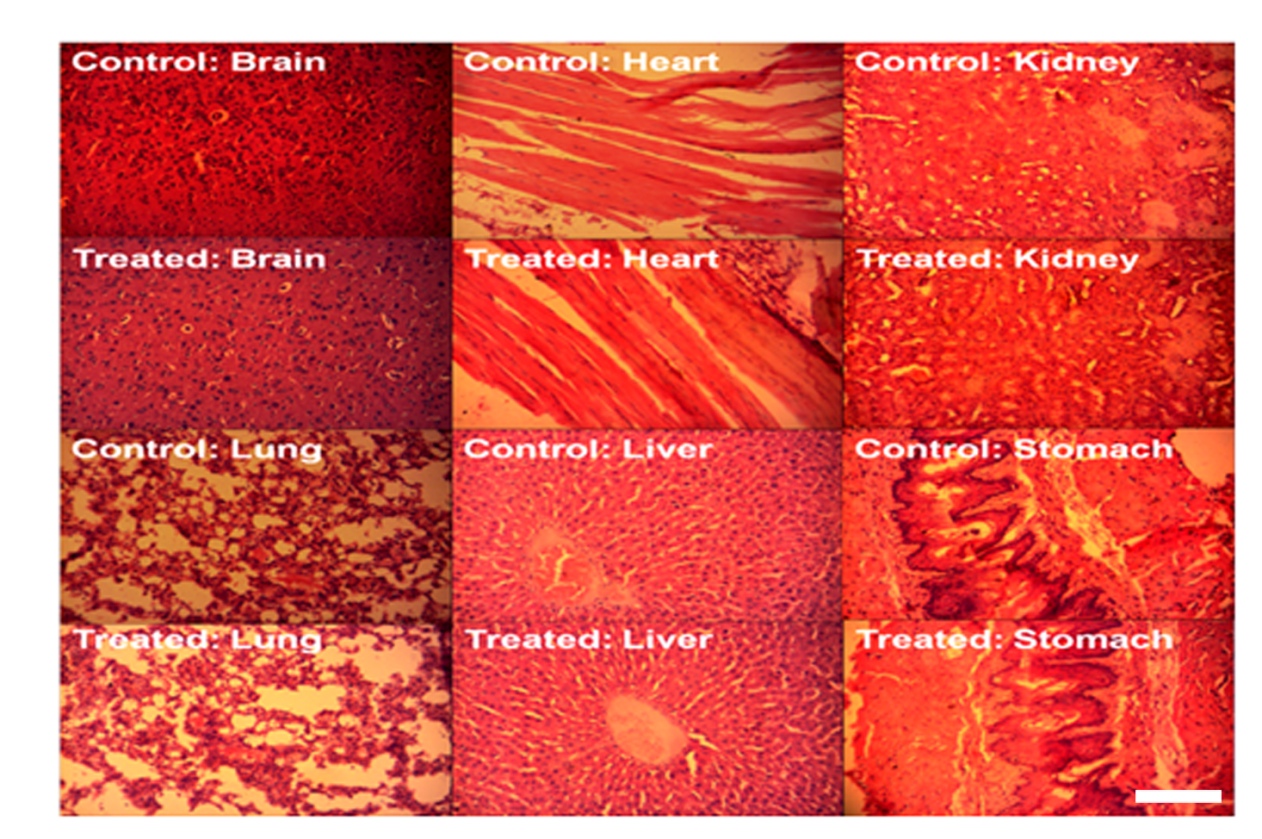

**Figure S3: Cellular architecture remains unchanged in BLEA-treated tumor-bearing mice** Fluorescent microscopic image showing histological sections of brain, lungs, heart, liver, kidney and stomach as indicated above. The toxicity evaluation of BLEA in different vital organs was visualized in hematoxylin and eosin stain (HE) (100X magnification). HE staining revealed no change in the cellular architecture of different tissues of control and treated groups. The scale bar represents 20 µm.

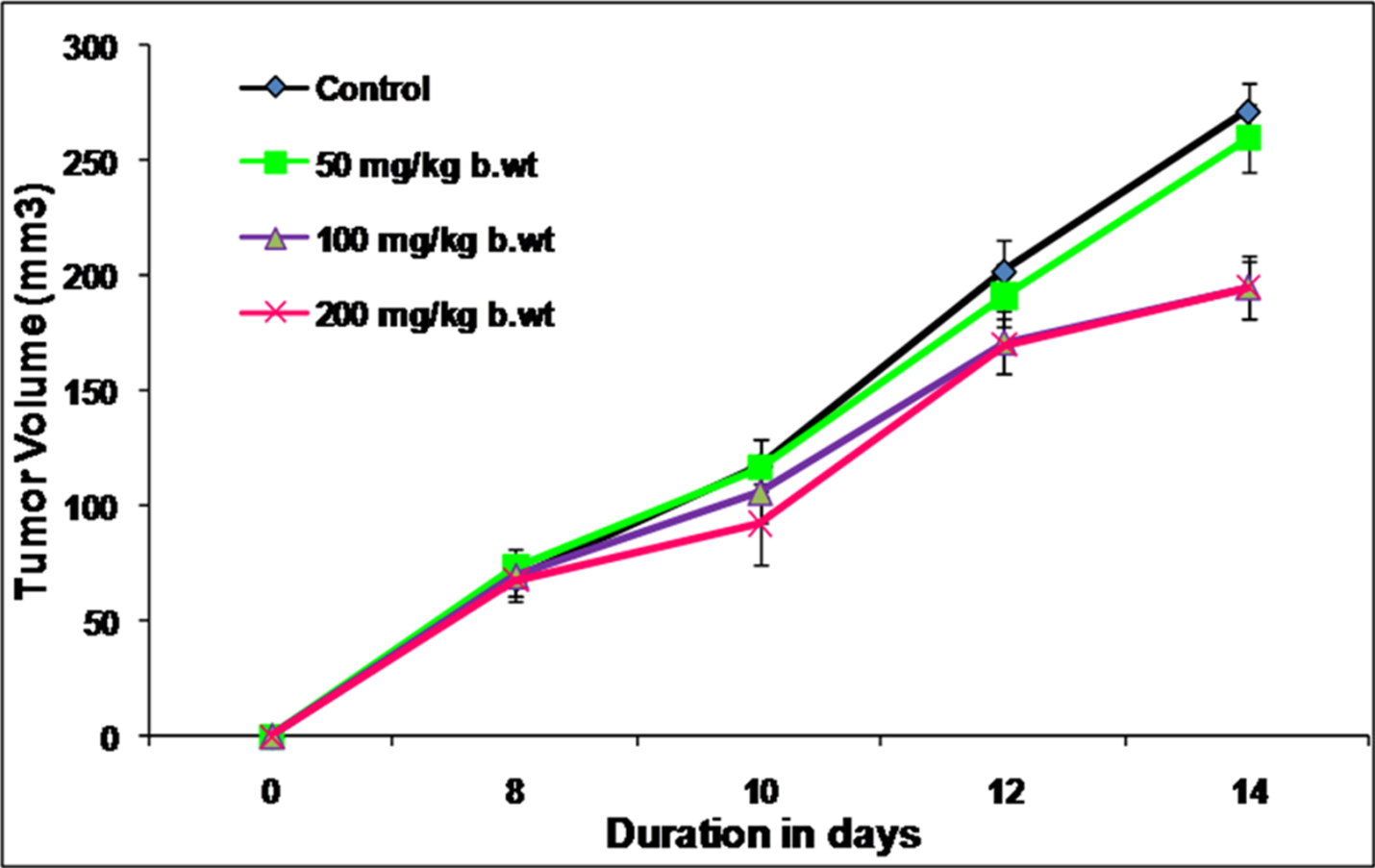

**Figure S4:** Determination of effective doses for BLEA. The graph shows the effectiveness of different doses of BLEA-derived leaf extract in mice. However, 200 mg/kg body weight of BLEA reduces tumor volume significantly in tumor-bearing mice compared to untreated control tumor mice.

**Supplemental Tables**

| Selective Index | MCF-7 | MDA-231 | HCT-15 | MIN6 | EAC | L929 |
| --- | --- | --- | --- | --- | --- | --- |
| BLPE | 14.92 | 5.07 | 4.69 | 24.61 | 3.79 | 1.00 |
| BLEA | 52.17 | 2.36 | 4.45 | 1.88 | 17.37 | 1.00 |
| BLM | 17.96 | 3.13 | 1.11 | 0.40 | 1.36 | 1.00 |

**Table S1:** The table shows the selective index of the different extracts against various cancer cell lines as indicated above.

| Observation | 0 h | | 0.5 h | | 1 h | | 2 h | | 4 h | | 24 h | | 14th day | |
| --- | --- | --- | --- | --- | --- | --- | --- | --- | --- | --- | --- | --- | --- | --- |
|  | D1 | D2 | D1 | D2 | D1 | D2 | D1 | D2 | D1 | D2 | D1 | D2 | D1 | D2 |
| Skin & fur | N | N | N | N | N | N | N | N | N | N | N | N | N | N |
| Mucous membrane | N | N | N | N | N | N | N | N | N | N | N | N | N | N |
| Tremors | N | N | N | N | N | N | N | N | N | N | N | N | N | N |
| Convulsion | N | N | N | N | N | N | N | N | N | N | N | N | N | N |
| Salivation | N | N | N | N | N | N | N | N | N | N | N | N | N | N |
| Diarrhea | N | N | N | N | N | N | N | N | N | N | N | N | N | N |
| Sleep | N | N | N | N | Y | Y | N | Y | N | N | N | N | N | N |
| Coma | N | N | N | N | N | N | N | N | N | N | N | N | N | N |
| Lethargy | N | N | N | N | N | N | N | N | N | N | N | N | N | N |
| Mortality | N | N | N | N | N | N | N | N | N | N | N | N | N | N |

**Table S2:** The table shows the observational studies of oral toxicity in BLEA-derived leaf extract in mice (OECD test 425).

| Compounds identified by GC-MS analysis | Mol. Wt. |
| --- | --- |
| Compound 1: 3-methoxy-2,5-dimethylpyrazine | 138.17 |
| Compound 2: 2,6-di-tert-butylcyclohexa-2,5-diene-1,4-dione | 220.31 |
| Compound 3: 4a-methoxy-1,1,2a,5-tetramethyldecahydro-1H-cyclopenta[cd]indene | 206.32 |
| Compound 4: (E)-2-methylhexadec-7-ene | 222.24 |
| Compound 5: icos-3-yne | 236.39 |
| Compound 6: trideca-1,12-diene | 238.45 |
| Compound 7: Palmitic acid | 278.52 |
| Compound 8: (E)-3,7,11,15-tetramethylhexadec-2-en-1-ol (Phytol) | 180.33 |
| Compound 9: (E)-octadec-9-enoic acid | 292.41 |
| Compound 10: (1-ethyl-4-methyl-1,1a,2,3,4,4a,9,10-octahydrocyclopropa[3',4']pyrido[2',3':3,4]cyclopenta[1,2-b]indol-10-yl)methyl acetate | 256.42 |
| Compound 11: 2-(((2-ethylhexyl)oxy)carbonyl)benzoic acid | 296.53 |
| Compound 12: 17-hydroxy-7,10,13,17-tetramethyl-1,2,6,7,8,9,10,11,12,13,14,15,16,17-tetradecahydro-3H-cyclopenta[a]phenanthren-3-one (Calusterone) | 282.46 |
| Compound 13: 3,5-dihydroxy-10,13-dimethyl-17-(6-methylheptan-2-yl)hexadecahydro-6H-cyclopenta[a]phenanthren-6-one | 338.44 |
| Compound 14: 10,13-dimethyl-17-(6-methylheptan-yl)hexadecahydrospiro[cyclopenta[a]phenanthrene-3,2'-[1,3]dioxolane | 278.34 |
| Compound 15: 2,4-di-tert-butylphenol | 316.48 |
| Compound 16: methyl3-(3,5-di-tert-butyl-4-hydroxyphenyl)propanoate | 418.65 |
| Compound 17: 2-phenyl-4H-chromen-4-one (Flavone) | 430.71 |

**Table S3:** The table shows the identification of various compounds of BLEA by GC-MS spectral analysis, as indicated above.
